# A Gene Expression Atlas of Lohmann White Chickens

**DOI:** 10.1101/2022.07.30.500160

**Authors:** Jiannan Zhang, Xinglong Wang, Can Lv, Yiping Wan, Xiao Zhang, Juan Li, Yajun Wang

**Affiliations:** Key Laboratory of Bio-resources and Eco-environment of Ministry of Education, Animal Disease Prevention and Food Safety Key Laboratory of Sichuan Province, College of Life Sciences, Sichuan University, Chengdu 610064, China

**Author notes:** Corresponding authors: Prof. Yajun Wang, Key Laboratory of Bio-resources and Eco-environment, Ministry of Education, College of Life Sciences, Sichuan University, Chengdu, 610065, China.; E-mail address Prof. Juan Li Key Laboratory of Bio-resources and Eco-environment, Ministry of Education, College of Life Sciences, Sichuan University, Chengdu, 610065, China.

**Keywords:** Transcriptome, Tissue expression profile, Annotation, GPCR, Sex-biased

## Abstract

Chicken (*Gallus gallus domesticus*) as one of the most economically important farm animals plays a major role in human food production and has been widely used as a key animal model that is presumed to be typical of avian and generally applicable to mammals in studies of developmental biology, virology, oncogenesis, and immunology. To get a better understanding of avian biology, global analysis of gene expression across multiple tissues is needed, which will aid genome annotation and support functional annotation of avian genes. We present a large-scale RNA-Seq dataset representing all the major organ systems from adult Lohmann White domesticus chickens. An open-access chicken tissue gene expression atlas (TGEA) (<u>chickenatlas.avianscu.com</u>) is presented based on the expression of 224 samples across 38 well-defined chicken tissues. Network-based cluster analysis of this dataset grouped genes according to dimensionality reduction and whole-body co-expression patterns, which were used to infer the function of uncharacterized genes from their co-expression with genes of known function. We describe the distribution and tissue specificity of 21,430 genes present in the chicken gene expression atlas and assign those signatures, where possible, to specific tissue populations or pathways. To better understand the functions of GPCRs in avian, we quantified the transcript levels of 254 nonodorant GPCRs in all tissues. Cluster analysis placed many GPCRs into expected anatomical and functional groups and predicted previously unidentified roles for less-studied receptors. We also produced this atlas to analyze male and female mRNA expression profiles in chicken somatic and gonad tissues. Our analyses uncovered numerous cases of somatic sex-biased mRNA expression, with the largest proportion found in the chicken pineal body, pituitary, and liver. This high-resolution gene expression atlas for chickens is, to our knowledge, the largest transcriptomic dataset of any avian to date. It provides a resource to improve the annotation of the current reference genome for chicken, presenting a model transcriptome for avian, and can be used as a resource for predicting roles for incompletely characterized GPCRs, exploring sex-biased specific gene expression, and for other purposes.

## Introduction

Domestic chickens are important livestock species that make significant contributions to the global agricultural economy in the form of eggs and meat (Mottet and Tempio, 2017). In addition, the domestic chicken (*Gallus gallus*) is widely used as a model for developmental biology. Compared to mammals, chickens are vertebrates that are phylogenetically distant from humans, but they play an important role in developmental biology because of their easy access to embryos (Davey and Tickle, 2007). The publication of the chicken genome (Red junglefowl) in 2004 paved the way for understanding the genetic basis for distinguishing the different phenotypes of domesticated chickens from their wild relatives (K. 1, 2004). Recent studies using population resequencing data and de novo assembled-based chicken pan-genome have improved the sequencing quality for the assembly of complete avian genomes and revealed that the numbers of chicken genes are comparable to those of other tetrapod vertebrates and a new pan-genome pattern of birds (Li et al., 2022; Wang et al., 2021). In addition to pan-genome analyses, transcriptome annotation is crucial for a wide array of biological research areas (Cunningham et al., 2022). The identification of complete libraries of transcriptional elements provides information about the functional roles and relationships of genomic motifs, which in turn can be compared to understand a wide range of biological mechanisms. However, due to the complexity of transcript splicing and the limitations of previous low-throughput technologies, the current state of chicken annotation with imprecise transcript models represents a typical example.

Microarray analysis has been widely used in many model species, including chickens (Burnside et al., 2005; Li et al., 2008). Microarrays are capable of studying hundreds of genes simultaneously and therefore help improve our understanding of the complexity of the chicken gene network. Although this technology offers great opportunities for the study of gene expression in a straightforward manner for a relatively low price, it is difficult to detect rare transcripts. The necessity of prior information on a large number of genes also limited the use of this technology for chicken. With the development and decreasing costs of NGS technologies, refers to technologies that allow for the concomitant sequencing of millions of smaller pieces of DNA, RNA-seq is rapidly supplanting microarrays for gene expression profiling in research (Conesa et al., 2016). When using RNA-seq to quantify the abundance of transcripts, information on the whole transcriptome can thus be acquired without prior knowledge of the transcripts in the target species (Wang et al., 2009). Analysis of RNA-Seq data can benefit from, but is not limited to, existing genomic knowledge and is well suited for non-model species that lack a high-quality reference genome.

For RNA-seq analysis in chickens, researchers have mostly used GRCg6a (or earlier versions), derived from a single red jungle fowl individual, as the reference genome in the past two decades, not the newly unpublished GRCg7b from a broiler. Nevertheless, it indicates that *G. gallus* domesticus is an admixed species, not only derived from red jungle fowl (Wang et al., 2020). A recent study also found different genome sizes between red jungle fowl and domestic chicken lineages (Piégu et al., 2020). This reference from red jungle fowl therefore cannot fully capture the genetic diversity of domesticated chickens and may be unable to reveal the genetic basis of some phenotypes. In addition to a high-quality pan-genome, a larger and more comprehensive high-resolution transcriptional map of different strains of domestic chickens is needed.

Atlases of gene expression are important tools for functional genomics. Groups of transcripts, members of which will have similar expression profiles, can be linked to a shared function, such as a specific pathway or biological process (Oliver, 2000). The tissue gene expression atlas (TGEA) has previously been used to annotate genes of unknown function in human (Andersson et al., 2014), pig (Freeman et al., 2012), and sheep (Clark et al., 2017). Transcriptomes of seven organs (cerebrum, cerebellum, heart, kidney, liver, ovary, and testis) across developmental time points from early organogenesis to adulthood for human, rhesus macaque, mouse, rat, rabbit, opossum and chicken were characterized to advance understanding of the genetic and developmental foundations of the evolution of mammalian phenotypes (Cardoso-Moreira et al., 2019). The co-expression information in these atlases also provides rich information about complex traits and disease susceptibility. Earlier studies in chickens mostly used red jungle fowl to construct the tissue gene expression atlas (TGEA) (McCarthy et al., 2019; Merkin et al., 2012). The first comprehensive transcriptome of the red jungle fowl was generated based upon RNA from 15 tissues collected from two red jungle fowls (one male and one female) (Merkin et al., 2012). Another tissue gene expression atlas (TGEA) derived from the assembly of short read RNAseq data of 20 tissue types from J-line layer chickens was created to compare and independently validate the PacBio transcriptome (Kuo et al., 2017). In addition, there is no other published large-scale tissue gene expression atlas available for domesticated chickens. A systematic global study of the tissue expression landscape in domesticated chickens is not available.

Domestic chickens are a significant source of animal protein all over the world. Various lines have been heavily selected for traits that will maximize production, such as increased egg production or quick weight gain. The chicken expression data from different breeds can provide support for understanding gene function, anatomy, and phenotype for agriculture and biomedical research. Previous studies suggest that sex-biased gene expression might be widespread and have potentially profound functional implications. Even so, previous efforts to identify sex-biased genes in mammals and birds were restricted to a single tissue, and many had limited resolution, because they were based on microarray technology and/or lacked biological replicates (Kaiser and Ellegren, 2006; Mank and Ellegren, 2009). To provide a more complete view of avian sex-biased genes, we have conducted the first comprehensive survey of male and female gene transcriptomes based on RNAseq of somatic and gonadal tissues from Lohmann White chickens. The Lohmann White Layer is one of the most successful commercial egg-laying breeds available, raised specifically for egg-laying productivity (Singh et al., 2009).

The gene expression atlas dataset of Lohmann White chickens presented here is the largest of its kind from any avian species to date and includes RNA-Seq libraries from tissues representing all the major organ systems of female and male chickens. Our aim was to provide a model transcriptome for domesticated chickens and give insight into gene and tissue function, the molecular basis of complex traits, underlying sex-biased gene expression patterns across tissues, and, using advanced visualization and clustering analysis techniques, we have identified networks of co-expressed genes. This data will support improved annotation of the chicken genomes and provide insight into biological processes underlying the complex traits that influence the productivity of chickens.

## Materials and Methods

### Chicken and sample collection

Adult chickens (Lohmann White Layer) used in this study were purchased from local commercial companies. A total of 8 chicken individuals were collected from sexually mature chickens (male: N = 4, female: N = 4) at the 1-year-old stage. All 224 samples of 38 tissues included in the chicken RNA Atlas are listed in **Supplementary Table 1**, including abdominal fat, adrenal gland, bursa of Fabritius, cecum, cerebellum, cerebrum, crop, duodenum, gizzard, heart, hindbrain, hypothalamus, ileum, infundibulum, jejunum, kidney, liver, lung, magnum, midbrain, muscle, ovary, pancreas, parathyroid glands, pineal body, pituitary, proventriculus, rectum, retina, skin, spinal cord, spleen, testis, thymus gland, thyroid gland, tongue, uterus, visceral fat. All tissue samples were stored at −80℃ before RNA extraction. All animal experiments were conducted in accordance with the Guidelines for Experimental Animals issued by the Ministry of Science and Technology of People’s Republic of China. All animal experimental protocols were approved by the Animal Ethics Committee of the College of Life Sciences, Sichuan University (Chengdu, China).

### Library preparation and sequencing

RNA extraction was performed with a RNAzol-based tissue RNA extraction protocol (Molecular Research Center, Cincinnati, OH). The tissue was homogenized mechanically with a pre-cooled glass homogenizer in liquid nitrogen. Total RNA was then extracted with a standardized protocol based on RNAzol reagent. Briefly, after precipitating with DEPC-treated ultra-pure water, the RNAzol lysed tissues were centrifuged (12,000 rpm, 10 min). BAN solution (4-bromoanisole) was added to purify the RNA and eliminate genomic DNA. Then, an equal volume of cold 100% isopropanol was added. The precipitated pellet was washed three times with 600 μL 75% ethanol and dissolved in 40 μL RNase-free water. The quality and quantity of total RNA samples were measured using a Onedrop1000 Spectrophotometer (Thermo Fisher Scientific, Waltham, MA, USA).

Library preparation was carried out using the DNBseq technology provided by MGI Tech Ltd. First, total mRNA and noncoding RNAs were enriched by removing ribosomal RNA (rRNA) using a MGIEasy rRNA depletion kit (MGI Tech, China). Enriched RNAs were then mixed with RNA fragmentation buffer resulting in short fragments. Third, complementary DNA (cDNA) was synthesized from the fragmented RNAs using N6 random primers, followed by end repair and ligation to BGIseq sequencer compatible adapters. The quality and quantity of the cDNA libraries were assessed using the Agilent 2100 BioAnalyzer (Agilent Technologies). Finally, the libraries were sequenced on the BGISEQ-500 with 150-bp paired-end read (PE150). An average of 41.36 million reads per sample were generated for each library. Sequencing reads that contained adapters, had low quality, or aligned to rRNA were filtered before following bioinformatics analysis.

### Raw data processing

For each tissue, a set of expression estimates, as transcripts per million (TPM), was performed using Salmon v.1.6.0 (Patro et al., 2017) and mapped to the NCBI chicken reference transcriptome (bGalGal1 maternal broiler GRCg7b), for the initial analysis. One of the novel and innovative features of Salmon is its ability to accurately quantify transcripts without having previously aligned the reads using its fast, built-in selective-alignment mapping algorithm. In order to ensure an accurate set of gene expression estimates, we employed a Salmon quasi-mapping-based mode by building a decoy-aware transcriptome index file. Salmon’s estimate of the number of reads mapping to each transcript was used for transcript-level abundance. Tximport v.1.24.0 (Soneson et al., 2015) was used to compute the gene-level estimates from transcript estimates. An overview of the total reads, Q30 clean reads, and mapping ratio is provided in **Supplementary Table 2**.

### Gene expression, network cluster analysis and annotation

Genes with less than 10 reads in all samples were excluded from further analysis. DESeq2 (Love et al., 2014) was used for DEGs detection and variance stabilizing transformation (vst) normalization. The output matrix of count data was normalized with a negative binomial distribution and values were log2 transformed. To identify sample-to-sample correlation, the Spearman’s correlation of DESeq2 normalized gene expression among tissues was computed. Statistical plots principal component analysis (PCA) and distance matrix analysis were generated with the same package to assess variance between sample groups and sample replicates. P-values were adjusted for multiple testing with the Benjamini and Hochberg method for controlling of the false discovery rate (FDR). Genes were called significant when the adjusted P-value was < 0.01. GO enrichment and KEGG pathway analyses were performed using the clusterProfiler software package on the R platform (Yu et al., 2012).

Graphia (Freeman et al., 2020) was used for network cluster analysis of the chicken tissues gene expression atlas. In brief, similarities between individual gene expression profiles were determined by calculating a Pearson correlation matrix for both gene-to-gene and sample-to-sample comparisons, and filtering to remove relationships where r < 0.80. A network graph was constructed by connecting the remaining nodes (genes) with edges (where the correlation exceeded the threshold value). This graph was interpreted by applying the Markov Cluster algorithm (MCL) at an inflation value (which determines cluster granularity) of 2.0. The local structure of the graph was then examined visually. Genes with robust co-expression patterns, implying related functions, clustered together, forming cliques of highly interconnected nodes (Clark et al., 2017). Expression profiles for each cluster were examined in detail to understand the significance of each cluster in the context of the biology of chicken tissues. Top Clusters (1 to 50) were assigned a functional “class” and “sub-class” manually by first determining if multiple genes within a cluster shared a similar biological function based on gene ontology, determined using the Bioconductor package “clusterProfiler”. The gene components of all clusters can be found in **Supplementary Table 3**.

### Data visualization

Data visualization was performed in R (v 4.2.1) using RStudio. The majority of visualizations were made using ggplot2 (v 3.3.6), supplemented with the following packages: pheatmap (v 1.0.12), ggpubr (v 0.4.0), tidygraph (v 1.2.0), figures were assembled, annotated, and aesthetically adjusted in Adobe Illustrator (Adobe Inc., San José, USA).

## Results and discussion

### Scope of the chicken tissue gene expression atlas dataset

The core of this dataset was derived from six one-year-old domesticated Lohmann White chickens. From these animals, we collected 38 tissues of all major organ systems and, to the extent possible, biological replicas of each sex (three female and three male), except for those of testes, bursa of Fabritius, adrenal gland (left), and oviducts, which represent three or four adult chickens. Some tissue samples, including hypothalamus, pituitary, adrenal gland (right), and liver, were supplemented by two additional animals of the same breed (*N* = 8). A detailed list of all tissues can be found in **Supplementary Table 1**. A wide range of tissues for the atlas were selected to obtain the largest diversity of transcripts possible. Tissues were chosen to give as comprehensive a set of organ systems as possible and to include those associated with phenotypic characteristics such as growth, metabolism, and reproduction.

### Sequencing depth and coverage

Approximately 9.266×10^9^ sequenced reads were generated from the TGEA libraries, with an average of 70–93% mapping rate per tissue. For each tissue, a set of expression estimates, as transcripts per million (TPM), were obtained using the fast and bias-aware quantification tool Salmon (Patro et al., 2017). Salmon has been developed that associates sequencing reads directly with transcripts without a separate quantification step. First, the RNA-seq data were log-transformed. After the normalization, a mean value from all replicates for each tissue separately was calculated. All genes that were not expressed in at least one tissue were removed from the analysis. The chicken reference genome (maternal broiler GRCg7b) includes 25,635 loci that are transcribed (18,023 protein coding), of which 21430 were detectable with expression of TPM >1, in at least one tissue from at least one individual, ranging from 8,442 to 16,229 genes detected per tissue type (**Supplementary Table 4**), in the final chicken gene expression atlas dataset, demonstrating the depth and scope of this dataset. Highly specialized tissue types, such as the pancreas, magnum, and proventriculus, express fewer genes, whereas tissues composed of many different cell types (e.g., testis, adrenal gland, pineal body, and anterior pituitary) express the highest number of genes, in line with results from human tissues (Djureinovic et al., 2014; Uhlén et al., 2015).

To further investigate similarities of global transcriptome profiles between tissues, Spearman correlation was used in a pairwise correlation heatmap for the 37 tissues. The heatmap with a sample-wide representation of all tissues shows that testis and the various brain samples have the most divergent global expression profiles, which is similar to the pattern in human and pig tissues (Karlsson et al., 2022; Uhlén et al., 2015). In general, closely clustering tissues often share germ layer origin, functions, and/or cellular composition. Our data also indicated the expression profiles of like tissues were correlated (Figure 1). The testes have a large enrichment of germ cells that are classified separately. Neuroectoderm-derived tissues such as brain tissues, pituitary gland, pineal gland, retina, and spinal cord cluster into one major region. Surprisingly, the adrenal gland tissues of domestic chickens had a high correlation with the above neuroectoderm-derived tissues. The gastrointestinal tract tissues, including rectum, cecum, duodenum, ileum, and jejunum, are clustered very closely.

**Figure 1.**
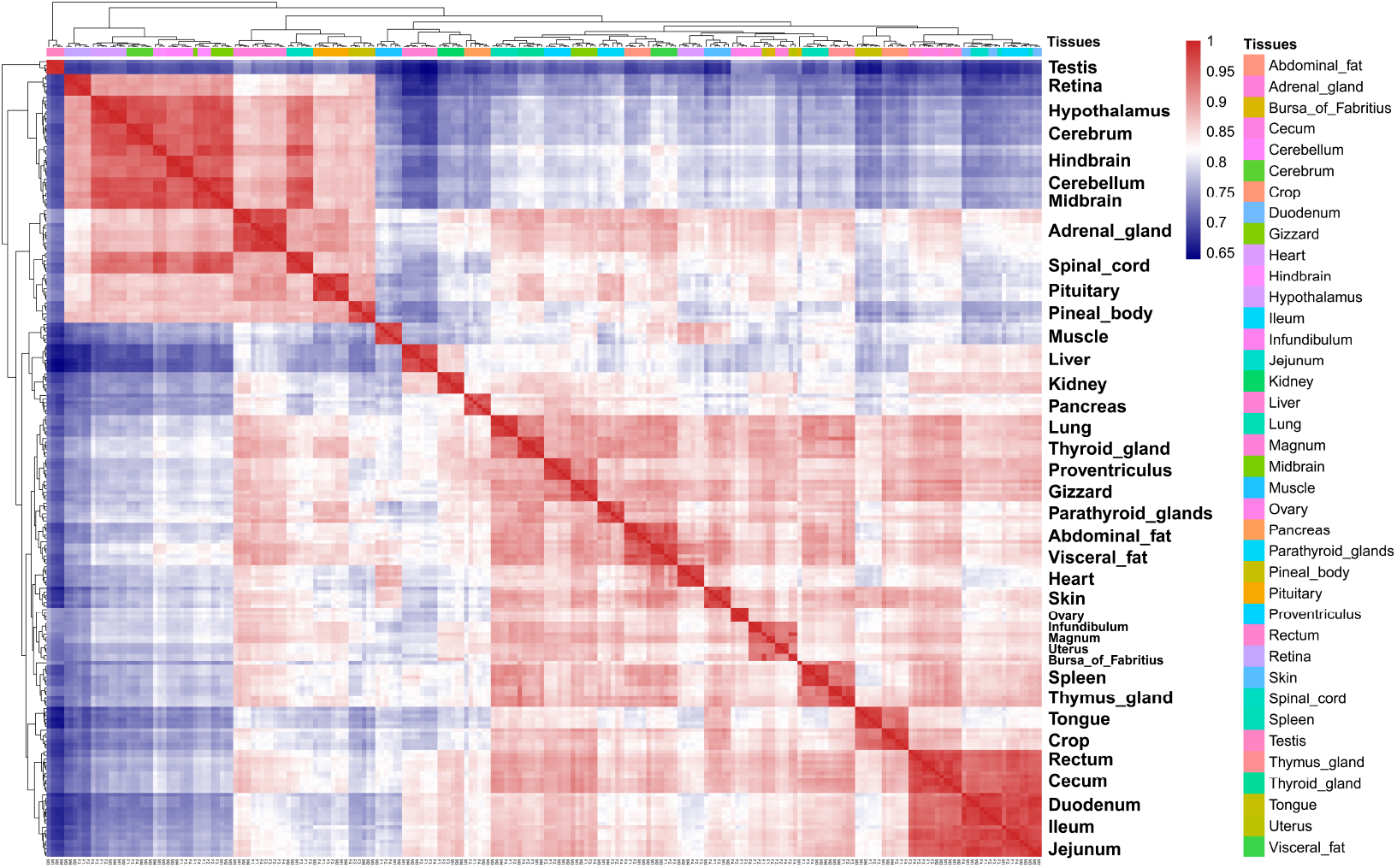
Heatmap showing the pairwise Spearman correlation of global expression across the 38 different tissue types of six chickens.

The genome-wide expression profiles were investigated for all the 350 individual samples using dimensional reduction analysis was performed to investigate the genome-wide expression profiles for all the 224 individual samples, and the results for principal component analysis (PCA) are shown in Figure 2. It shows that tissue types with related functions share similar global expression profiles and that the brain and small and large intestine samples have a unique expression pattern compared to other tissues. The 30 brain subregions cluster according to the basic organization of the brain, with the spinal cord separated from the pituitary and pineal body. Taken together, these results validate the aggregation of data from multiple tissues to create an informative expression atlas.

**Figure 2.**
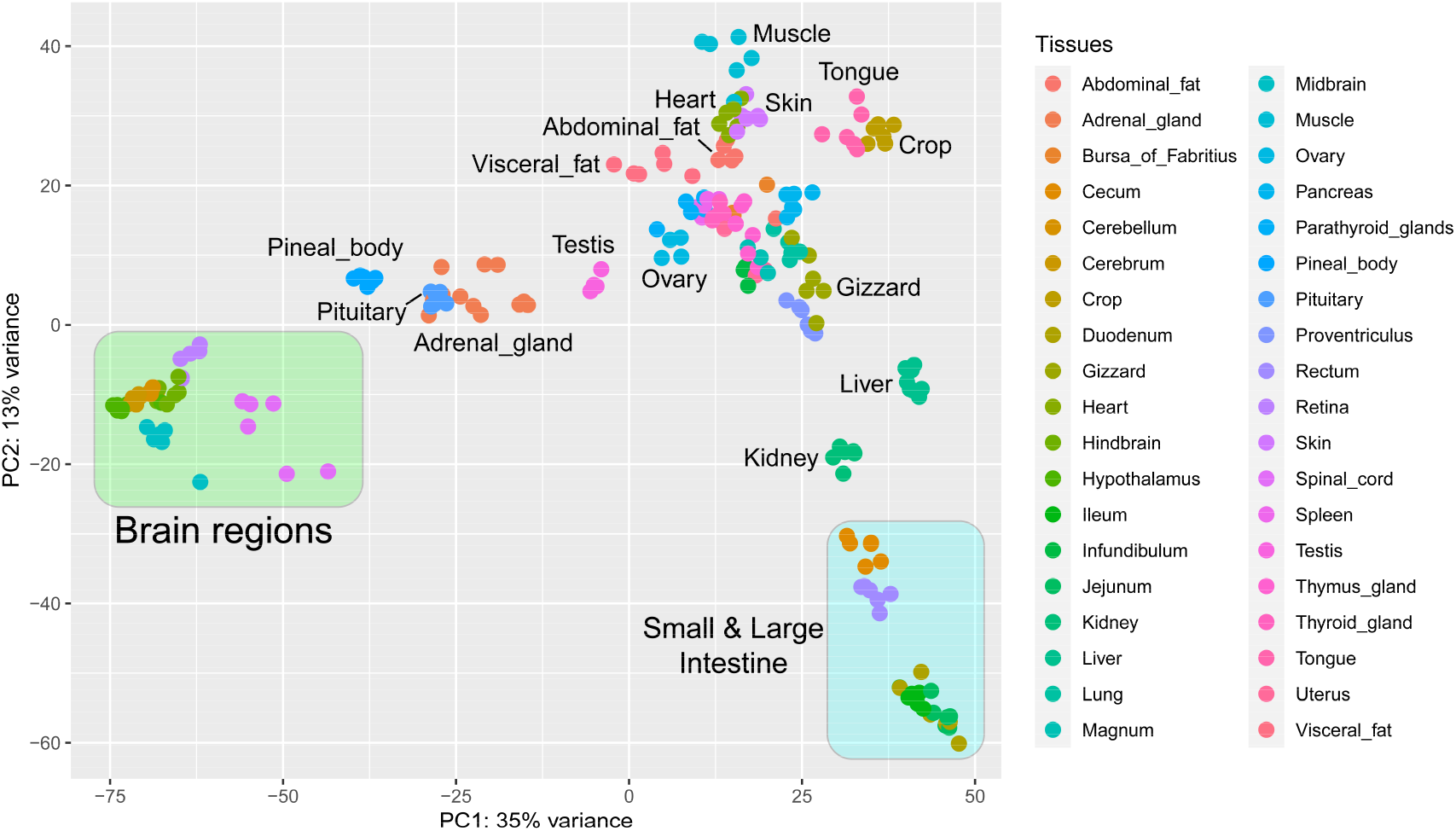
Principle component analysis (PCA) plot showing the relation and clustering of all chicken tissue samples.

### Network cluster analysis and Data visualizing

Network cluster analysis of the chicken tissue gene expression atlas was generated using Graphia Enterprise, a tool for the visualization and analysis of network graphs from big data, including protein interaction data, transcriptomics, and single cell analyses (Dimonaco et al., 2020; Freeman et al., 2020). Graphia filters out low and stably expressed genes, thereby highlighting the most variable and potentially tissue-specific genes. A gene-to-gene pairwise Pearson correlation matrix across all samples was created with a Pearson correlation co-efficient threshold of r = 0.80 and an MCL (Markov Cluster Algorithm) inflation value of 2.0 to identify clusters of co-expressed genes. To remove noise, genes with an average TPM >10 in at least one tissue and unannotated genes were restricted in the analysis. The gene-to-gene network comprised 14,308 nodes (transcripts) and 717,150 edges (correlations above the threshold value). Clusters are numbered according to their relative sizes, with the largest cluster being cluster 1, and so on. Clusters with fewer than five nodes were excluded from further analysis, resulting in 24 components ranging from 5 to 4,040 nodes. The network graph is shown in Figure 3A, along with the expression profiles of selected clusters. The graph consisted of one large component containing 4,040 nodes, and 23 smaller components. Genes found in each cluster are listed in **Supplementary Table 3**, in which clusters are labelled according to the tissue showing the highest expression in the cluster.

**Figure 3.**
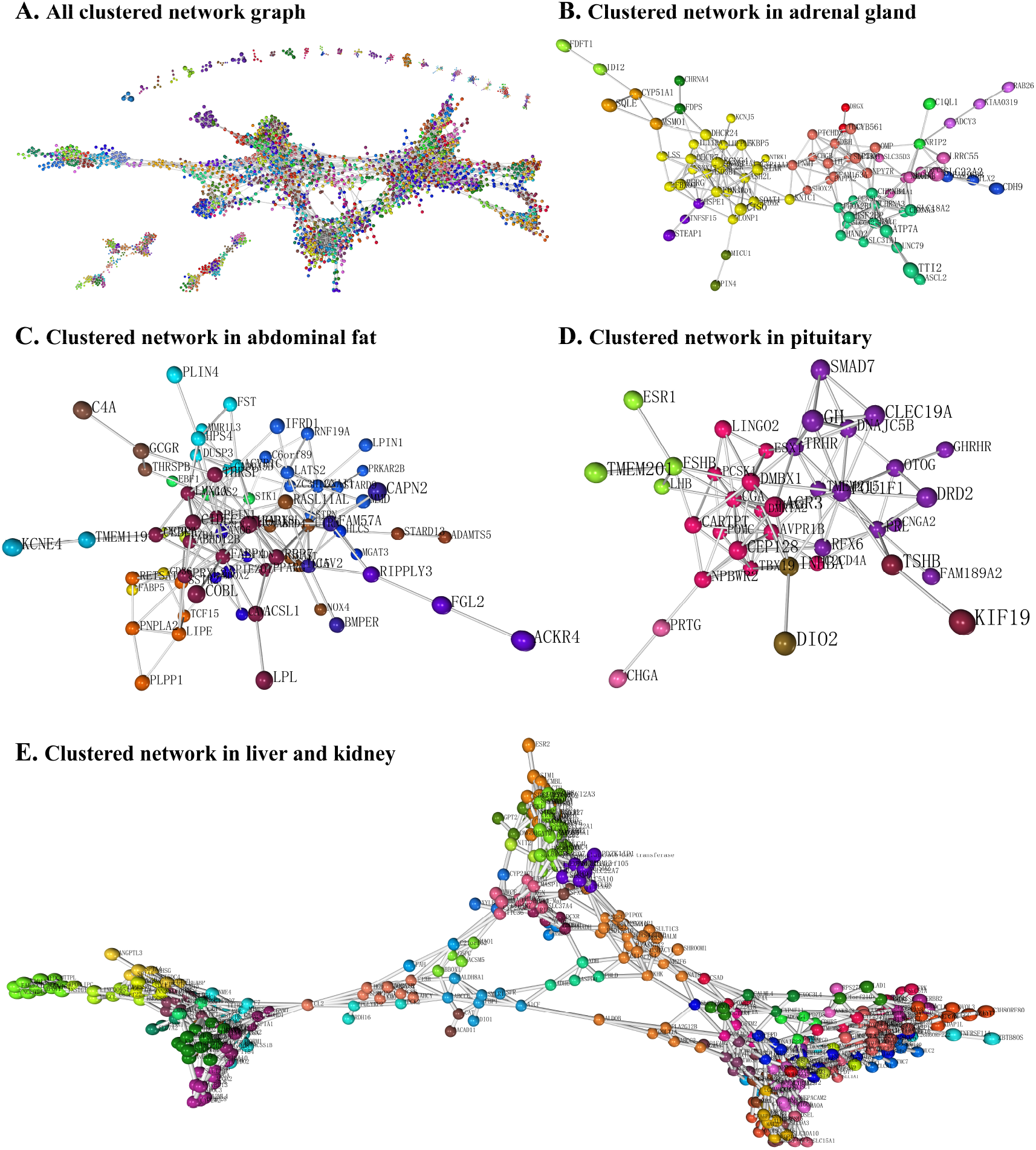
Network visualization and clustering of the chicken gene expression atlas. (**A)** A three-dimensional visualization of a Pearson correlation gene-to-gene graph of expression levels derived from RNA-Seq data from analysis of all chicken tissues. (**B-D**) The tissue specificity of gene co-expression in three selected clusters are shown, including adrenal gland (**B**), abdominal fat (**C**), pituitary (**D**), and liver/kidney (**E**). Each node (sphere) in the graph represents a gene and the edges (lines) correspond to correlations between individual measurements above the defined threshold. Co-expressed genes form highly connected complex clusters within the graph. Genes were assigned to groups according to their level of co-expression using the MCL algorithm.

The tissue-specific expression patterns observed across clusters are highly similar to those observed in pigs, sheep, humans and mice (Clark et al., 2017; Freeman et al., 2012; Su et al., 2002, 2004). Most co-expression clusters included genes exhibiting a specific tissue expression pattern (Figure 3A). There were some exceptions, including the largest cluster (cluster one), which contains ubiquitously expressed “housekeeping” genes, encoding proteins that may play roles in all cell types. With a few exceptions, the remaining co-expression clusters were composed of genes exhibiting expression only in a distinct tissue, such as the adrenal gland (component 6) (Figure 3B), abdominal fat (component 7) (Figure 3C), or pituitary (component 10) (Figure 3D).

Some co-expression is shared between two or more organ systems and is associated with known shared functions. For example, the second largest component 2, with 481 nodes (Figure 3E), exhibiting high expression in the liver and kidney, is enriched by KEGG for expression of genes relating to the biosynthesis of cofactors, carbon metabolism, glycolysis/gluconeogenesis process, and PPAR signaling pathway. It contains many genes that encode enzymes involved in amino acid biosynthesis (e.g., *PRPS2*, *PSPH*, *CTH*, *MAT2L*, *ACY1*, *SDSL*, *MAT1A*, *PAH*, *GPT2*, and *ALDOB*) and tryptophan metabolism (*MAOA*, *HAAO*, *KMO*, *ALDH8A1*, *KYNU*, *TDO2*, *EHHADH*, *IDO2*, *CAT*, and *KYAT1*). The contribution of the kidney and liver to amino acid metabolism is well known in humans (Stumvoll et al., 1998) and rodents (Ayyar et al., 2017). These observations suggest that shared catabolic pathways in the liver and kidney are largely conserved in chicken, and detailed collation of genes in this cluster could provide further specific insights. In addition, we identified a large number of genes specifically expressed in abdominal adipose (Figure 3C) and pituitary tissues (Figure 3D).

### The adrenal gland

Stringent co-expression clustering requires that each transcript be quantified in a sufficiently large number of different states to establish a strong correlation with all other transcripts with which it shares coordinated transcription and, by implication, a shared function or pathway. The impact of this approach, which was effective for profiling the expression of region-specific genes in specific tissue, was evident from the pig and sheep gene expression atlases (Clark et al., 2017; Freeman et al., 2012). The paired chicken adrenal glands are located anterior and medial to the cephalic lobes of the kidneys (Scanes and Dridi, 2021). In this study, we sampled and sequenced the adrenal tissues from both sides separately to identify genes that might be associated with adrenocortical function in development, maturation, and stress. We identified three main adrenal clusters (clusters 14, 64, and 91) from the atlas data (Figure 4A). Genes in cluster 14 showed the highest expression levels in adrenal glands with many of the genes encoding well-characterized steroidogenesis-specific proteins, including *STAR*, *CYP11A1*, *CYP21A1*, *HSD3B1*, *HSPD1*, *LSS*, *DHCR7*, *DHCR24*, and *SOAT1*. Genes in this cluster were enriched for KEGG pathways including “steroid biosynthesis” (p = 1.91 × 10^-11^), “aldosterone synthesis and secretion” (p = 7.10 × 10^-6^), and “cortisol synthesis and secretion” (p = 1.46 × 10^-5^) (Figure 4B).

**Figure 4.**
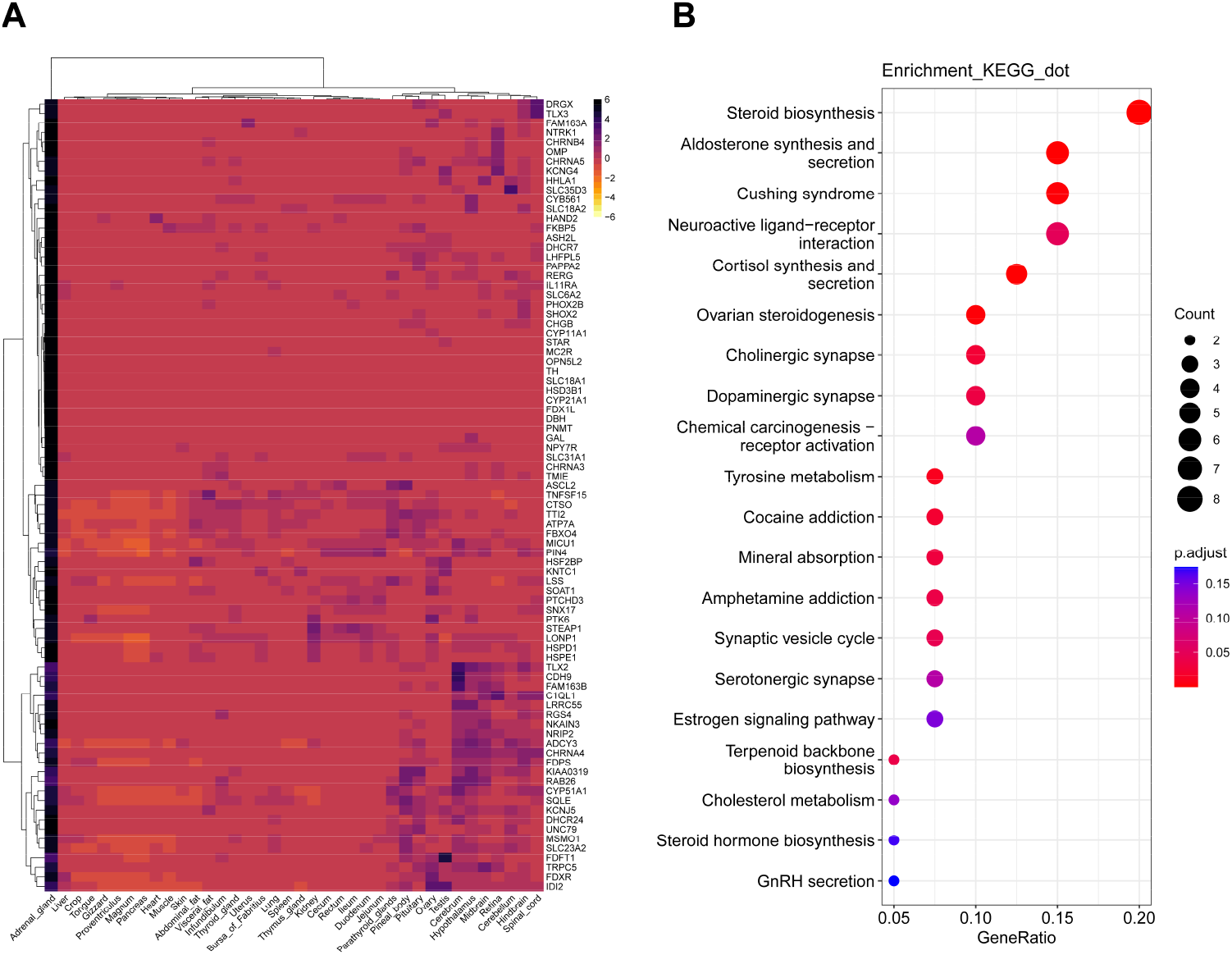
Heat-map (**A**) showing the mRNA levels of the genes in adrenal gland clusters, which were enriched for KEGG pathway as shown by dotplot (**B**).

The expression of *STAR* and *CYP11A1* in chicken adrenal glands differs from human, mouse, pig, and sheep, and is restricted to adrenal glands with specific high expression, in which the expression level of *STAR* was nearly 500-fold higher than that of ovarian tissue (Karlsson et al., 2022). Adrenal steroidogenesis is carried out by microsomal and mitochondrial enzymes, which are highly conserved among vertebrates (Castillo et al., 2015). STAR is the hormone-sensitive, steroidogenic acute regulatory protein that delivers free cholesterol from the outer mitochondrial membrane to the inner mitochondrial membrane, which is primarily localized to the gonads and adrenals, known as the “classical” steroidogenic organs, in mammals (Anuka et al., 2013; Bauer et al., 2000). In agreement with our findings, relatively high expression of *STAR* was also observed in the hen adrenal, instead of gonads (Bauer et al., 2000). Among chicken ovarian tissues, the highest level of *STAR* expression was observed in the granulosa layer of the F1 follicle (Bauer et al., 2000). The expression pattern of *CYP11A1* in domestic chickens is also significantly different from that of mammals. The *CYP11A1* gene encodes the cholesterol side-chain cleavage enzyme, also termed cytochrome P450scc, which catalyzes the conversion of cholesterol to pregnenolone in the first step of steroid biosynthesis in mitochondria. It is expressed in the mammals adrenals and gonads under the control of pituitary peptide hormones (Guo et al., 2007), but our findings showed that *CYP11A1* was more abundant in adrenals (∼8000 TPM) than in gonad tissue (∼240 TPM). These results suggest that the adrenals may contribute precursors to gonadal steroidogenesis in chickens. *CYP11A1* expression levels was found to have similar expression levels in zebrafinch testis, ovary and adrenal (Freking et al., 2000). It is indicated that the expression pattern of steroidogenesis-specific genes may not be uniform among the avian species in that in some species there may be an “adrenalgonadal unit” in which the adrenals contribute precursors for gonadal steroidogenesis.

In addition, those genes, including *LSS*, *DHCR7*, and *DHCR24* involved in cholesterol biosynthesis, were highly expressed in the liver and adrenal glands of mammals (Karlsson et al., 2022). By comparison, the TPM level of chicken *LSS*, *DHCR7*, and *DHCR24* showed peak expression in the adrenal glands, which was nearly 10-fold higher than that of liver tissue. In mammals, cholesterol is either absorbed from dietary sources or synthesized de novo. The liver and intestinal mucosa are the main sites of cholesterol synthesis. Up to 70–80% of cholesterol in humans is synthesized de novo by the liver, and another 10% is synthesized de novo by the small intestine (Yang et al., 2020). The *LSS* gene encodes lanosterol synthase and a null mutation for lss decreased cholesterol levels in rats (Mori et al., 2006). DHCR7 is the enzyme that catalyzes the reduction of 7-dehydrocholesterol to cholesterol in the last step of cholesterol biosynthesis (Horling et al., 2012). DHCR24 is the enzyme that catalyzes the cholesterol biosynthesis by reducing the delta-24 double bond of desmosterol (Cecchi et al., 2008). Previous studies have demonstrated that three enzymes are key factors contributing to the hydrogenation of dehydrocholesterol. Interestingly, our mapping shows that the adrenal gland appears to contain the key enzymes required for local de novo cholesterol synthesis, which may be a potentially important tissue for cholesterol synthesis in chickens.

### Anatomical Profiling of G Protein-Coupled Receptor Expression

G protein-coupled receptors (GPCRs) are the most commonly exploited targets in modern medicine. The GPCR superfamily comprises the largest and most diverse group of surface receptors in mammals (Regard et al., 2008). GPCRs are differentially expressed throughout the organism, respond to diverse endogenous ligands, and regulate a host of physiological processes, including hemostasis, immune function, metabolism, neurotransmission, and reproduction (Lagerström et al., 2006). Although in vivo roles have been defined for many of the GPCRs in mice and humans, the expression and function of many such receptors are incompletely characterized in avian species, including chickens, and a significant fraction remain orphans in vertebrates (Kooistra et al., 2021; Vassilatis et al., 2003). Relying on our established domestic chicken TGEA database, we analyzed the pattern of GPCR mRNA expression across tissues and the relative abundance of each of 254 nonodorant GPCRs in 38 tissues from adult chickens. Hierarchical clustering analysis revealed groupings of tissues and receptors that predicted physiological functions for individual receptors and receptor clusters (Figure 5A). The resulting dendrograms for both the tissue and receptor axes showed functional clusters.

**Figure 5.**
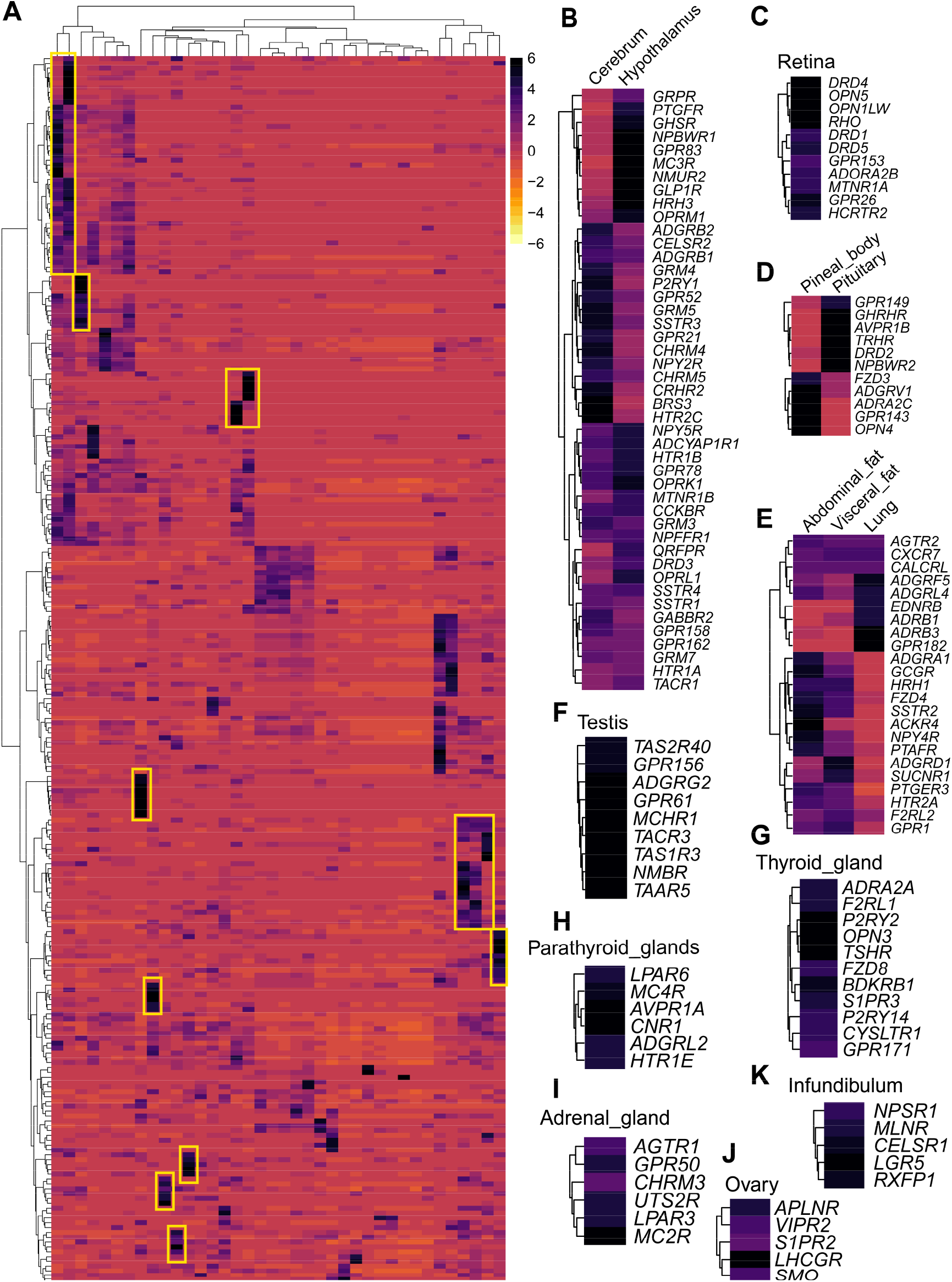
Hierarchical clustering of GPCR expression across chicken tissues. (**A**) Transformed RNA-Seq data for the 254 GPCRs assayed in the 38 tissues was evaluated by unsupervised hierarchical clustering with average linkage with R package pheatmap. Multiple clusters and subclusters were seen, and ten were chosen for further analysis. (**B**) A small portion of the “CNS” cluster, by far the largest. This portion contains receptors for important neurotransmitters including serotonin, neuropeptide Y, orexin and opiates. (**C**) The retinal cluster contains light-sensing opsins as well as other receptors known to regulate vision. (**D**) The “pituitary-pineal body” cluster contains many well-documented regulators of pituitary and pineal body function, including *GHRHR*, *TRHR*, *OPN4*. (**E**) A portion of the “adipose” cluster is shown; note the abundance of *GCGR*, *SSTR2*, *NPY4R*, *PTAFR* and *FZD4*. (**F-J**) unique groups of receptors clusters in testis (**F**), thyroid gland (**H**), adrenal gland (**I**), ovary (**J**), and infundibulum (**K**) were shown.

The CNS tissues cerebrum, hypothalamus, retina, cerebellum, spinal cord, hindbrain, and midbrain clustered together, as did the immune/hematopoietic tissues spleen and thymus gland. The steroidogenic organs adrenal gland and ovary, clustered with the parathyroid glands, infundibulum, uterus, Bursa of Fabritius, and skin. However, the testes had a very distinct pattern of GPCR expression. Liver, kidney, heart, muscle, gizzard, crop, tongue, pancreas, magnum, and proventriculus formed a group, as did large ileum, duodenum, jejunum, cecum, and rectum. Pituitary and pineal body formed a cluster, perhaps related to their common endocrine function. Abdominal fat and visceral fat formed an “adipose” cluster. The lung and thyroid gland formed a cluster, perhaps related to development from the endoderm.

A number of receptor axis clusters were easily recognized (Figure 5). The CNS cluster was by far the largest. Our analysis, as well as others (Regard et al., 2008; Vassilatis et al., 2003), suggests that more than 80% of all nonodorant GPCRs are expressed in the CNS. A small portion of the “CNS” cluster in the cerebrum and hypothalamus is shown in Figure 5B. Included are *GPRP*, *PTGFR*, *GHSR*, *MC3R*, *NMUR2*, *ADGRB2*, *GPR52*, *SSTR3*, *GRP21*, *NPY2R*, *CRHR2*, *BRS3*, *HTR2C*, *NPY5R*, *HTR1B*, *GPR78*, *CCKBR*, *NPFFR1*, *DRD3*, and *HTR1A*, all of which have been implicated in the regulation of neuronal function in vertebrates (Kooistra et al., 2021).

The highly expressed genes in retina tissue were shown in Figure 5C, which contains *DRD4*, *OPN5*, *OPN1LW*, *RHO*, *DRD1*, *DRD5*, *GPR153*, *ADORPA2B*, *MTNR1A*, *GPR26*, and *HCRTR2*, all of which are known to function in the eye (Chen and Palczewski, 2016; Chen et al., 2016). The pituitary cluster (Figure 5D) includes *GPR149*, *GHRHR*, *AVPR1B*, *TRHR*, *DRD2*, and *NPBWR2*. Correspondingly, *FZD3*, *ADGRV1*, *ADRA2C*, *GPR143*, and *OPN4* were highly expressed in the pineal body. It is widely assumed that hypothalamic GHRH activates GHRH receptor (GHRHR) to stimulate GH synthesis and release in the pituitary of chickens (Wang et al., 2007). Interestingly, NPBWR2 has recently been previously implicated in pituitary gland function as a novel inhibitory secretagogue for GH, prolactin, and ACTH in chickens (Bu et al., 2016; Liu et al., 2022). In this study, DRD2 was demonstrated to mediate the effect of dopamine to inhibit VIP-induced cPRL expression in the chicken anterior pituitary (Lv et al., 2018). Chicken TRHR1 plays important roles in mediating TRH actions on the chicken pituitary, such as involvement in the regulation of TSH, GH, and PRL secretion (Harvey, 1990; Harvey et al., 1978; Scanes, 1974). AVPR1B was thought to partially mediate the AVT-induced cPOMC/cPRL expression in anterior pituitary (Wu et al., 2019). In mammals, *Gpr149* was highly expressed in the ovary and also in the brain, and *Gpr149* knockout mice displayed increased fertility and enhanced ovulation (Edson et al., 2010; Karlsson et al., 2022). Considering the specific high expression of *GPR149* in the pituitary gland of domestic chickens, the question of whether GPR149 signaling in chickens play a role similar to that found in mammals awaits further investigation.

Finally, unique groups of receptor clusters in the testis, thyroid gland, adrenal gland, infundibulum, and ovary were shown in Figure 5F-5J. The expression and function of most receptors in these groups have been widely studied, but many unique expression features that have not been reported were found in this TGEA database. For example, testes showed a GPCR expression pattern very distinct from that of other tissues. *TAS2R40*, *GPR156*, *ADGRG2*, *GPR61*, *MCHR1*, *TACR3*, *TAS1R3*, *NMBR*, and *TAAR5* were almost perfectly specific to testes. It is unknown whether such receptors play a role in spermatogenesis or other testicular functions and their potential utility as targets for drugs aimed at controlling fertility. Whether these receptors play a role in spermatogenesis or other testicular functions in chickens is unknown.

Taken together, the results outlined above reveal that tissues cluster into largely expected functional groups purely on the basis of their GPCR repertoires, and both expected and unique groups of receptors cluster by tissue function. These data identify sets of receptors involved in specific aspects of physiology and should prove useful in providing clues regarding in vivo roles for orphan GPCRs and new roles for receptors with known ligands in avian.

### Sex-biased gene expression in chickens

Sex-specific differences in gene expression have been reported in humans (Imahara et al., 2005), mice (Everhardt Queen et al., 2016), cattle (Forde et al., 2016) and pigs (Zhang et al., 2013). DESeq2 was used to examine male and female biased gene expression in chicken tissues (except for non-somatic tissues such as the testis, ovary, infundibulum, magnum, and uterus) (Love et al., 2014). For all somatic tissues and each tissue, we used DESeq2 to identify mRNAs with sex-biased expression at a global false discovery rate (FDR) of 5%. Differential expression between testis and ovary tissues (gonads) was analyzed and corrected separately due to the mixture of sex-related and tissue-related effects, which otherwise reduced the stringency of the significance test for the somatic samples. An overview of sex-biased gene expression for all somatic tissues and single tissues is given in Figure 6, with complete details available in **Supplementary Table 5**. We identified several individual genes that were sex-biased in multiple somatic tissues from a single species (**Supplementary Table 6**).

**Figure 6.**
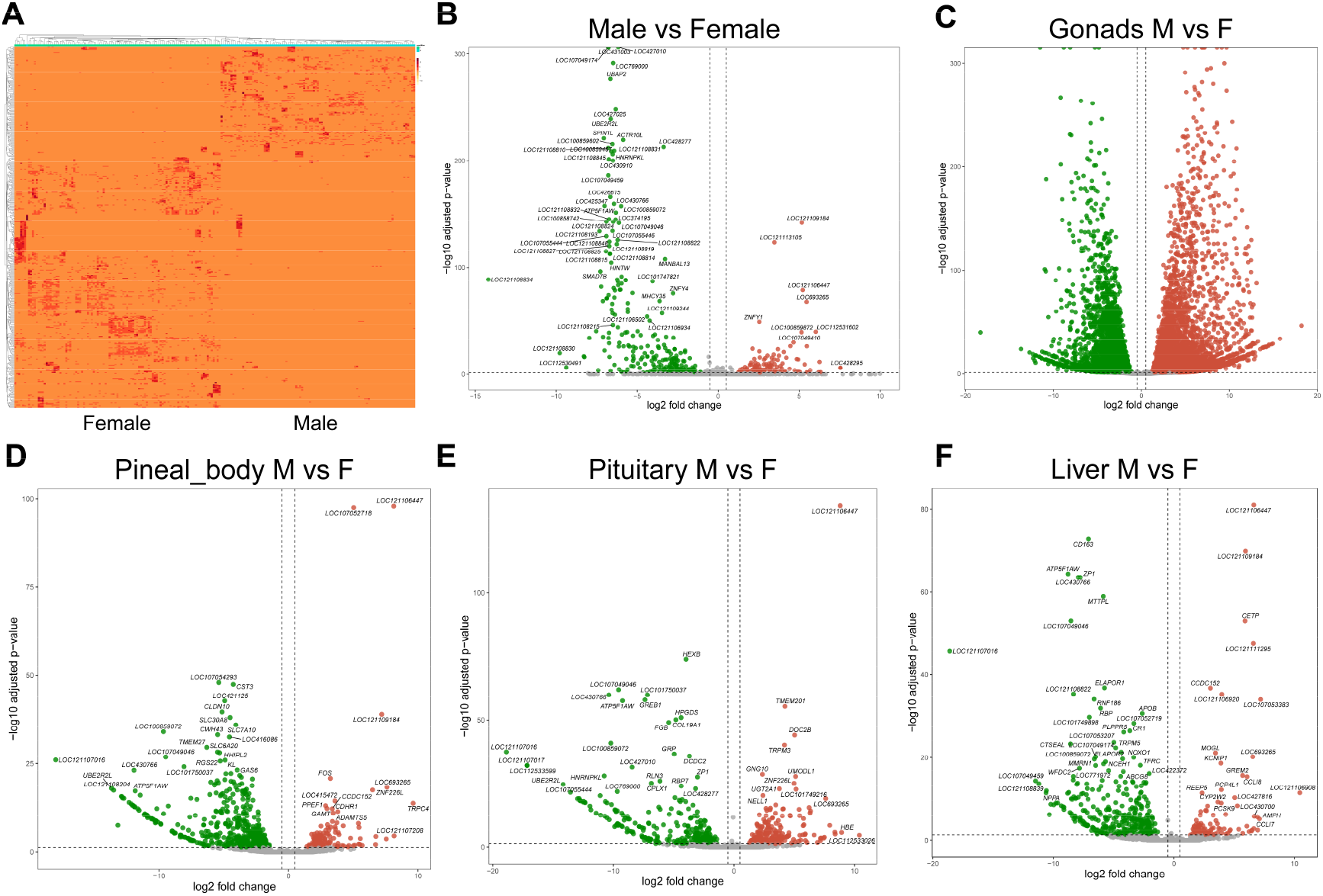
Sex-biased genes expression in chicken. Different expression genes (DEGs) were calculated for three male and three female replicates per tissue and the significance assessed with DESeq2, performed separately for somatic and gonadal samples. (**A**) Two-dimensional hierarchical clustering of different expression genes in 32 somatic tissues of six chickens. A set of 478 most differently active genes are clustered on the vertical axis, with individual tissues clustered on the horizontal axis. (**B**) Volcano plots showing the distribution of log2 gene expression of somatic tissues in female and male chickens (values > 0: female-biased genes; values < 0 male-biased genes) and significance levels. Red/green color marks significant genes (padj < 0.05). (**C**) Volcano plots showing that in addition to the sex-biased miRNA expression found in somatic tissues, we consistently found pronounced sex differences in the gonads. (**D-F**) Among the somatic tissues, the largest proportion was found in chicken pineal body (**D**), pituitary (**E**) and liver (**F**).

We found evidence of sex-biased differentially expressed genes (DEGs) in all of the investigated 32 somatic tissues, with a total proportion in whole somatic tissues (478 DEGs out of 22292 genes) (Figure 6A-6B), In addition to the sex-biased miRNA expression found in somatic tissues, we consistently found pronounced sex differences in the gonads (Figure 6C). Among the somatic tissues, the largest proportion was found in chicken pineal body (659 of 18124 genes) (Figure 6D), pituitary (530 of 18435 genes) (Figure 6E) and liver (468 of 15457) (Figure 6F), with the smallest proportion found in chicken tongue (80 of 16296), pancreas (88 of 13753), and proventriculus (105 of 14584). These findings are consistent with the large numbers of protein-coding genes that are expressed in a sex-biased manner in various tissues of the mouse (Yang et al., 2006). It is indicated that most sex-biased genes in somatic tissues are autosomal. The expression pattern of these genes can be explained through direct or indirect regulation by hormones or other sex-biased factors.

The top 15 female-biased DEGs (with well annotation in NCBI database, and basemean > 10) in somatic tissues are *SMAD7B*, *ST8SIA3L*, *SPIN1L*, *UBAP2*, *UBE2R2L*, *HINTW*, *HNRNPKL*, *ATP5F1AW*, *ACTR10L*, *THEM4*, *MTTPL*, *MHCY15*, *ZP1*, *YLEC8*, and *YLEC13*, and the top 15 male-biased DEGs are *ZNF226L*, *CCLI7*, *CCDC152*, *HCK*, *ZNFY1*, *OZFL*, *HIST1H2B8*, *NAA38*, *MHCY32*, *CETP*, *CENPK*, *PMM2*, *TMEM70*, *MSH3*, and *SNAPC5*. Among these genes, some candidate genes were identified as potential regulators for establishing sexual identity. *SMAD7B*, which is conserved in the SMAD family, was cloned from the developing chicken limbs in a previous study (Vargesson and Laufer, 2009). However, the functions of SMAD7B have not been determined. Conversely, histidine triad nucleotide-binding protein W (*HINTW*) is a well-analyzed W-linked candidate gene characterized by strong expression patterns in female embryos (Hori et al., 2000). Gene amplification and conversion in *HINTW* coincide with this being the only gene on the avian W chromosome so far found to have evolved a distinct female-specific function, which should be particularly beneficial to genes that have specialized in function subsequent to sex chromosome differentiation (Backström et al., 2005). In agreement with previous studies, *ZP1* was highly expressed in the female chicken liver, which was indicated to be synthesized in the liver and transported via the bloodstream to the follicle (Bausek et al., 2000, 2004). *CCDC152*, located on the W-chromosome, was highly expressed in the male kidney and liver, which was identified to reduce cell proliferation and migration through the JAK2/STAT signaling pathway in chicken (Lin et al., 2017). a *PMM2* also had higher expression in male chickens, which was found in metabolic pathways such as amino sugar and nucleotide sugar metabolism and fructose and mannose metabolism (Kanehisa and Goto, 2000). Differential regulation of PMM2 has been associated with virus infection of chicken embryo fibroblast cell cultures (Maślikowski et al., 2010). We also found that male chickens exhibited significantly higher levels of autophagy related 10 (*ATG10*) mRNA in the testis and kidney, which is consistent with previous reports (Piekarski et al., 2014). *ATG10* is a critical gene for autophagy and cancer, and there is increasing evidence for the importance of autophagy-related genes in the maintenance, therapy, and pathogenesis of cancer (Arya et al., 2018). Some of the sex differentially expressed genes obtained in this study have been analyzed in previous studies, but we found more unreported sex differentially expressed genes, which provide clues for the subsequent in-depth analysis of avian sex differences. In addition, several candidates that may regulate the expression of other genes were identified in this study. In the future, functional analysis of sex-biased genes in chicken is required.

### Sexually dimorphic gene expression in chicken anterior pituitary

In the chicken pituitary, some genes with a high TPM value (>200) are expressed in a sex-dependent manner, including *LOC121107016*, *LOC121107017*, *LOC112533599*, *CPLX1*, *RLN3*, *GRP*, *HPGDS*, *HEXB*, *SPARCL1* in female, and *HSPA5*, *PLK2*, *CSRP2*, *CCDC80*, *LOC112532140*, TMEM201 in male (Figure 6E). Interestingly, in our previous scRNA-seq study, we found that in gonadotrophs clusters of the chicken anterior pituitary, some genes (*CPLX1*, *RLN3*, *GRP* and *HPGDS*) are also expressed in a sex-dependent manner (Zhang et al., 2021). *RLN3* is highly expressed in female pituitary gonadotrophs but not in male pituitary gonadotrophs. Similarly, *GRP*, *HPGDS*, and *CPLX1* have relatively higher mRNA levels in female than that in male gonadotrophs, which was confirmed by using quantitative real-time PCR (qPCR) to indicate the sexually dimorphic expression patterns of *GRP*, *RLN3*, and *HPGDS* at the whole anterior pituitary level (Zhang et al., 2021). In addition, some novel differentially expressed genes were revealed in this study. *SPARCL1*, which has been proposed to mediate skeleton mineralization in these vertebrates (Venkatesh et al., 2014), was found to be highly expressed in the female pituitary. *CCDC80*, expressed highly in male pituitary and fat tissues, is a crucial regulator of energy homeostasis *in vivo* and functions as an inhibitor of adipogenesis (Grill et al., 2017). The sexually dimorphic expression pattern of genes in pituitary tissues may have important physiological relevance. For example, *GRP* and *RLN3*, which are highly expressed in female (but not male) pituitary gonadotropins, may play important roles in female reproduction. Future studies of these sex-biased genes in chickens and other birds will help reveal not only their effects on growth, metabolism, stress, and reproduction in males and females, but also their association with phenotypic characteristics of birds (e.g., egg-laying performance, reproductive mode, and meat production).

### Visualization of the expression atlas

We have provided the chicken tissue gene expression atlas (TGEA) as a searchable database hosted on the website (https://chickenatlas.avianscu.com/) via the Shiny Server Open Source software (Chang et al., 2017). The Shiny application allows users to view and download the expression profile of any given gene across tissues. The data was tabulated using the dplyr package and plotted using the ggplot2 package (Conesa et al., 2016). An R Shiny application to interact with the database was developed using the following R packages: shiny, shinyfeedback, and shinycssloaders (Sali and Attali, 2017). An example profile of the melanocortin receptor accessory protein 2 (*MRAP2*) gene from chicken is included in Figure 7, which is highly consistent with the tissue distribution we reported previously (Zhang et al., 2017). The chicken TGEA expression profiles are based on TPM estimates from the alignment-free Salmon output for the Lohmann White libraries, averaged across samples for ease of visualization. It is important to note that there may be a degree of variation in the expression patterns of specific genes between individuals, which was displayed using box plots. In addition, gene names and gene descriptions et al. of each gene were listed on the website to allow comparison between species, which is not the case for all genes in the atlas due to the limit of gene annotation in the current database. In parallel to the alignment-free Salmon method, a similar expression matrix was obtained by using an alignment-based approach to RNA-Seq processing with the HISAT, StringTie, and Ballgown pipeline (Pertea et al., 2016).

**Figure 7.**
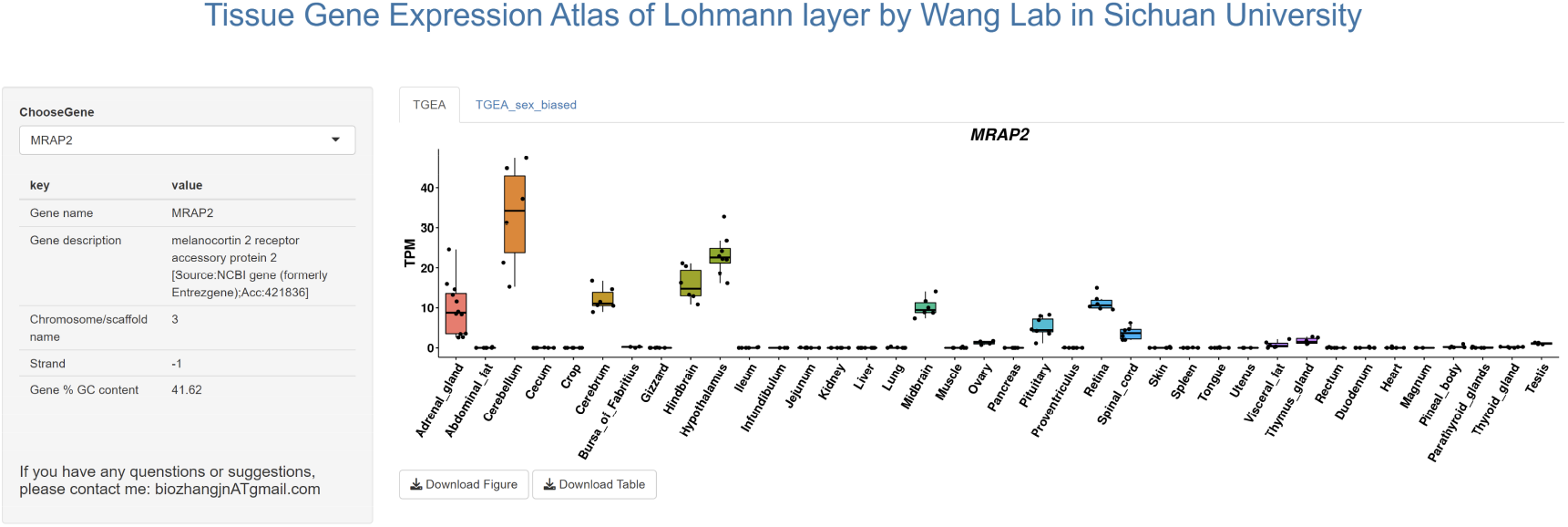
Screenshot of the expression profile of the chicken melanocortin receptor accessory protein 2 (*MRAP2*) gene within the Lohmann White chicken online tissue gene expression atlas (TGEA). Expression estimates from the chicken TGEA are available via the Shiny Server. This provides a searchable database of genes, with expression profiles across tissues for each gene displayed as histograms via the following link, https://chickenatlas.avianscu.com/. The Shiny Server platform supports searching for genes and downloading the high-solution figures, allows access to the raw data, and links to external resources.

## Conclusions

In conclusion, we present a transcriptional landscape of the chicken, covering 38 tissues and organs, to allow annotation of chicken genes based on an expression analysis, providing the largest gene expression dataset from an avian species to date. A genome-wide resource of the transcriptome map across all major tissues in chicken has been launched, and the data is available as an open-access resource called the chicken TGEA Atlas (chickenatlas.avianscu.com) with the gene expression profile across all tissues, including a comparison to the human and pig orthologs. Using the network clustering approach, we have been able to recapitulate known expression and functional relationships between genes as well as infer new ones based on guilt by association. The detailed analysis of the transcriptional landscape of genes encoding GPCRs showed that they highly exhibited tissue-specific patterns of expression and were enriched for distinct functions and pathways. This resource will aid studies to understand GPCR function and may assist in the identification of therapeutic targets in vertebrates. The TGEA atlas also provided evidence of the global expression of the sexually dimorphic genes, which is helping to understand the important pathways and networks with differential expression levels in males and females. Above all, this resource will facilitate future attempts to understand chicken biology and to use chicken as an animal model system for biomedical research, physiological, morphological, behavioral, and other aspects of development.

## Availability of data and materials

The data for the current study are available from the corresponding author upon reasonable request.

## Ethics statement

All animal experiments were conducted in accordance with the Guidelines for Experimental Animals issued by the Ministry of Science and Technology of People’s Republic of China. All animal experimental protocols were approved by the Animal Ethics Committee of the College of Life Sciences, Sichuan University (Chengdu, China).

